# Identification of Alpha-Gal conjugated Lipids in Saliva of Lone-Star Tick (*Amblyomma americanum*)

**DOI:** 10.1101/2024.02.22.581476

**Authors:** Surendra Raj Sharma, Sabir Hussain, Shailesh K. Choudhary, Scott P. Commins, Shahid Karim

## Abstract

Alpha-Gal Syndrome (AGS) is a delayed allergic reaction triggered by IgE antibodies targeting galactose-α-1,3-galactose (α-gal), prevalent in red meat. Its global significance has increased, with over 450,000 estimated cases in the United States alone. AGS is linked to tick bites, causing sensitization and elevated α-gal specific IgE levels. However, the precise mechanisms and tick intrinsic factors contributing to AGS development post-tick bites remain unclear. This study aims to characterize the alphagal conjugated lipid antigens in *Amblyomma americanum* (*Am. americanum*) salivary glands and saliva. Nanospray ionization mass spectrometry (NSI-MS) analysis revealed the identification of α-gal bound lipid antigens in *Am. americanum* saliva. Additionally, activation of basophils by isolated alpha-gal bound lipids and proteins provides evidence of their antigenic capabilities.

## Introduction

Alpha-gal syndrome (AGS) is a distinctive allergic response triggered by galactose-α-1,3-galactose (alpha-gal, α-gal), a disaccharide moiety on the end of glycan found in mammalian meat but notably absent in catarrhine primates, including humans (Commins et al., 2009; Macher et al., 2008). The lack of α-gal in humans is attributed to the inactivation of the α1,3-Galactosyl transferase (α1,3GT) gene in an ancestral Old-World species, setting humans apart from most other mammals (Macher et al., 2008). The presence of α-gal sugar is significant as it triggers the production of anti-Gal antibodies, including immunoglobulin M, immunoglobulin A, and immunoglobulin G in humans (Macher et al., 2008; Galili, 1999). In a more specific context, AGS originates from a unique immune response characterized by specific immunoglobulin E (sIgE) antibodies, specifically targeting α-gal antigens. This immune reaction leads to allergic responses typically becoming apparent 2-6 hours after the consumption of red meat or its derivatives (Commins et al., 2009;2011; 2014; Fischer et al., 2014). The key initiator of AGS is an α-gal antigen, which is frequently found in mammalian glycoconjugates like glycolipids and glycoproteins (Commins et al., 2009; Apostolovic et al., 2014; Fischer et al., 2014; Hamsten et al., 2013; Iweala et al., 2020; Macher et al., 2008). The formation of these glycoconjugates within organisms relies on a diverse family of enzymes known as glycosyltransferases (Berg et al., 2014; Roseman, 2001). These enzymes are predominantly present in the cells, tissues, and fluids of mammals, with the notable exception of humans, apes, and old-world monkeys (Apostolovic et al., 2014; Galili & Avila, 1999; Iweala et al., 2020; Takahashi et al., 2014). Consequently, the deactivation of α1,3GT in humans is thought to be the fundamental cause of allergy development when glycoconjugates containing alpha-gal antigens are orally ingested (Commins et al., 2014; Sharma and Karim, 2021).

Ticks are ectoparasites with the ability to transmit various disease-causing agents, macromolecules, and other substances to humans (Adegoke et al., 2020; Bullard et al., 2019; Chmelař et al., 2016). Numerous studies provide compelling evidence establishing a connection between tick bites and the emergence of red meat allergy, commonly known as alpha-gal syndrome (Commins et al., 2011; Binder et al,2021, Sharma and Karim 2021). In a previous study, we reported the presence of the α-gal antigen in the salivary gland extracts and saliva of *Am. americanum* and *Ixodes scapularis* using N-glycome profiling and proteome analysis (Crispell et al., 2019). Our follow-up studies provided compelling evidence that repeated exposure to tick salivary gland extract and tick bites results in a significant increase in antibody response against α-gal, inducing AGS in an alpha-gal knock out murine model (Choudhary et al., 2021; Sharma et al., 2024).

While the role of N-glycan alpha-gal antigens in ticks and the contribution of tick bites to AGS induction have been well-established (Crispell et al., 2019; Sharma et al., 2024), a critical aspect concerning the involvement of lipid-bound alpha-gal in AGS induction remains to be investigated. Recent scientific reports have suggested lipid antigens may be crucial in allergic sensitization (Roman-Carrasco et al., 2021; Iweala et al., 2021, Hopkins et al., 2022). However, a fundamental knowledge gap exists regarding whether ticks possess α-gal bound lipid antigens in their saliva or salivary glands and whether these antigens can sensitize the host. If present, how is the tick-related machinery linked to the development of lipid-bound alpha-gal antigens following hematophagy? Therefore, this study aims to identify α-gal conjugated lipid antigen markers in tick salivary glands and saliva.

## Materials and Methods

### Ethics statement

Animal experiments were performed adhering to the NIH Guideline for the Care and Utilization of Laboratory Animals. Tick feeding on sheep was approved by the University of Southern Mississippi’s IACUC under protocol #15101501.2, with measures to minimize animal discomfort.

### Ticks and other animals

Unfed adult Lone-star ticks *Am. americanum* were obtained from Ectoservices, USA, and received care at the University of Southern Mississippi in adherence to a standardized protocol (Patrick & Hair, 1975). Before infesting ticks on a sheep, the adult ticks were kept at room temperature with approximately 90% humidity, following a photoperiod of 14 hours of light and 10 hours of darkness. Depending on the specific experimental design, adult ticks were permitted to feed on sheep for durations ranging from 1 to 11 days to facilitate subsequent tissue collection.

### Tick tissue dissection and salivary gland and saliva extraction

Partially engorged female ticks were pulled off the sheep and underwent dissection, with subsequent removal and cleansing of their salivary glands and midguts in an ice-cold M199 buffer. The salivary glands of partially fed ticks (3-7 days post infestation, dpi) were dissected, pooled and flash frozen in liquid nitrogen and stored at –80°C until used. Tick saliva was obtained by prompting partially-blood-fed (4-7 dpi) female *Am. america*num to salivate into capillary tubes, employing the modified pilocarpine induction method as outlined in previous study (Crispell et al., 2019). The collected saliva was flash frozen in liquid nitrogen and promptly stored at −80°C for the future use.

### Lipid extraction, saponification, purification, permethylation, and analysis by nanospray ionization MS/MS

A total of 200 μl of pooled tick saliva (isolated from 50 partially feed ticks (3-7 dpi) and pooled salivary glands (from 40 pairs of SG) were obtained from partially fed *Am. americanum* ticks and promptly flash-frozen using liquid nitrogen and shipped to UGA (Complex Carbohydrate Research Center) in dry ice for subsequent analysis. The salivary glands were disrupted while maintaining cold conditions, in the presence of 50% MeOH. The resulting mixture was then transferred to a glass tube containing an additional 50% MeOH, adjusting the final composition to 4/8/3 (by volume) of Chloroform: Methanol: Water. Subsequently, the solution underwent a 3-minute sonication and was then centrifuged to separate the components. The supernatant was carefully transferred to a clean glass tube and subjected to drying. This process was repeated three more times, and all collected supernatants were combined into the same tube. The supernatants were evaporated with nitrogen before undergoing saponification, which involved the removal of phosphoglycerolipids. A methanol-based alkaline solution was added to the sample, and the lipids were allowed to incubate at 37°C overnight. The reaction was then neutralized by adding an equal volume of acetic acid while cooling on ice. To further purify the sample, salts were removed through tC18 SPE extraction. The samples were washed with water and subsequently eluted with methanol before undergoing drying once more. Free fatty acids were eliminated from the sample by introducing hexane.

For improved stability, sequencing via MS/MS, and enhanced sensitivity in mass spectrometry analysis, the isolated glycosphingolipids (GSLs) underwent permethylation. Briefly, the GSLs were dissolved in dimethyl sulfoxide and methylated with NaOH and methyl iodide. The reaction was stopped with water, and the per-O-methylated carbohydrates were extracted using methylene chloride, followed by drying under nitrogen. The samples were then analyzed using direct infusion nanospray ionization mass spectrometry (NSI-MS) with MS/MS fragmentation to confirm the structural composition. For analysis, the samples were reconstituted in a mixture of 2:1 MeOH: H2O with 1 mM NaOH and introduced to the mass spectrometer at a flow rate of 1 μL/min. FlexAnalysis (Bruker Daltonics) was used for data processing.

### Basophil Activation Test

Peripheral blood mononuclear cells (PBMCs) were sourced from a healthy donor with no α-gal allergies (α-gal sIgE <0.10) and isolated through a Ficoll–Paque (GE Healthcare in Chicago, IL, USA) gradient. Within the PBMC fraction, IgE was stripped from basophil using a cold lactic acid buffer (containing 13.4 mM lactic acid, 140 mM NaCl, and 5 mM KCl) for 15 minutes. Subsequently, basophils were sensitized overnight with plasma from α-gal allergic individual in RPMI1640 cell culture media (Corning CellGro in Manassas, VA, USA) containing 1 ng/mL IL-3 (R&D Systems in Minneapolis, MN, USA) at 37°C with 5% CO2.

Following sensitization, PBMCs were stimulated for 30 minutes with either basophil activation media alone (RPMI1640 containing 2 ng/mL IL-3), or test reagents in basophil media including rabbit anti-human IgE (1 μg; obtained from Bethyl Laboratories Inc. in Montgomery, TX, USA), lipid isolated from tick saliva, partially fed salivary gland and remaining protein fractions of fed salivary gland and saliva from *Am. americanum* (50 μg). Stimulation reactions were halted with 20 mM EDTA, and PBMCs were stained with fluorescently labeled antibodies targeting CD123 (from BioLegend in San Diego, CA, USA), human lineage 1 markers (CD3, CD14, CD16, CD19, CD20, CD56, from BD Biosciences in San Jose, CA, USA), HLA-DR, CD63 (from eBiosciences ThermoFisher in Waltham, MA, USA), and CD203c (from IOTest Beckman Coulter in Marseille, France). Staining was performed in a flow cytometry staining buffer containing 2% FBS and 0.02% NaN3. The samples were analyzed using an Attune NxT flow cytometer (Thermo Fisher Scientific, Waltham, MA USA, and data were processed using FlowJo v10.9.0 software from FlowJo LLC in Ashland, OR, USA. Statistical analysis utilized Prism version 7.03 from GraphPad Software in La Jolla, CA, USA, employing Mann– Whitney U-tests to compare the frequency of CD63+ basophils observed after stimulation with various compounds, with statistical significance set at a p-value of < 0.05.

## Results

### Mass-spectrometry reveals the presence of α-gal bound antigens in tick saliva

Glycan-bound lipid profiling of partially fed *Am. americanum* tick saliva and partially fed salivary gland using Nanospray ionization mass spectrometry (NSI-MS) with MS/MS fragmentation revealed critical information regarding the presence of alpha-gal bound lipids in tick. For the first time, we report that the saliva of partially fed *Am. americanum* tick contained various glycoforms possibly belonging to the isoglobo GB3 series possessing alpha-gal bound glycosylation (Table 1, Figure 1). In *Am. americanum* tick saliva the overall abundance of alpha-gal bound lipid was 28.8 percent of the total glycan-bound lipids. More specifically, NSI-MS revealed that partially fed *Am. americanum* saliva consisted of three different glycoforms with an observed mass of 1010.7, 1036.6, 1094.84, 1120.86,1214.85; 1298.98; 1324.92 along with an overall abundance of 35.5%, 15.5%,1.4%,18.9%,17.3%; 0.1% and 11.4% respectively (Figure 1; Table 1). In addition, NSI-MS also revealed the presence of four other ceramides in higher abundance (∼52% in total) (Table 1). We also analyzed lipids isolated from tick salivary gland structures revealing the presence of various glycoforms with an observed mass 1038.7450, 1066.77, 1092.82, 1311.90, 1337.94, 1557.03, 1583.07,20813.3, 2109.3, 2135.4 with relative abundance of 6%, 30.6%, 33.1%, 2.0%, 1.4%, 3.6%, 1.9%,2.7%, 13.3% and 5.3 % respectively (Supplementary table 1S). Indeed, salivary gland profile showed that high diversity of lipids possibly possessing 18,22,24 carbon with one or no unsaturation, belonging to ceramide (size range 1038-3040, supplementary table 1). Glycan signature belonging to these lipids lacked actual alpha-gal epitope however contained glycan chain of either glucose, mannose and GlcNAc in, glucose, mannose. In addition, some other additional lipids (size 1557.03 and 1583.07, supplementary tables1 potentially having 18,22 or 24 carbon with one or no-unsaturation were reported to be possessing glycans containing chain of GalNac, GulNAc, glucose and mannose. Furthermore, lipids found in saliva with higher mass 2081.3, 2109.3 and 2135.4 had similar core glycosylated pattern containing gal, GalNaC,GlcNAc linked to disaccharides chain of mannose and glucose (Supplementary table 1S, Supplementary figure 2S-4S).

**Table 1:**
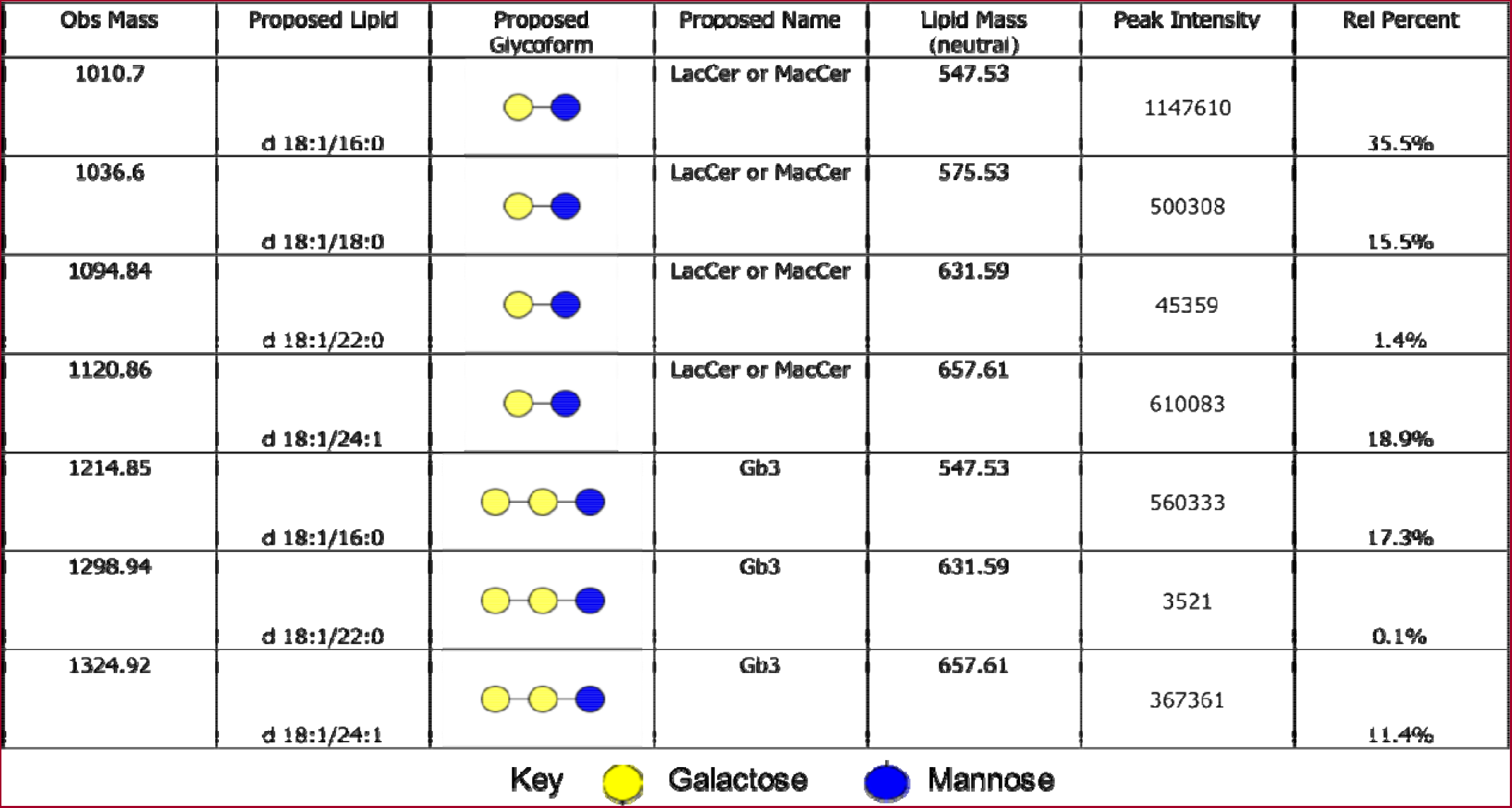
Relative percentages of each glycolipid observed in tick saliva.

| Obs Mass | Proposed Lipid | Proposed Glycoform | Proposed Name | Lipid Mass (neutral) | Peak Intensity | Rel Percent |
| --- | --- | --- | --- | --- | --- | --- |
| 1010.7 | d 18:1/16:0 |  | LacCer or MacCer | 547.53 | 1147610 | 35.5% |
| 1036.6 | d 18:1/18:0 |  | LacCer or MacCer | 575.53 | 500308 | 15.5% |
| 1094.84 | d 18:1/22:0 |  | LacCer or MacCer | 631.59 | 45359 | 1.4% |
| 1120.86 | d 18:1/24:1 |  | LacCer or MacCer | 657.61 | 610083 | 18.9% |
| 1214.85 | d 18:1/16:0 |  | Gb3 | 547.53 | 560333 | 17.3% |
| 1298.94 | d 18:1/22:0 |  | Gb3 | 631.59 | 3521 | 0.1% |
| 1324.92 | d 18:1/24:1 |  | Gb3 | 657.61 | 367361 | 11.4% |

Key Galactose Mannose

**Figure 1:**
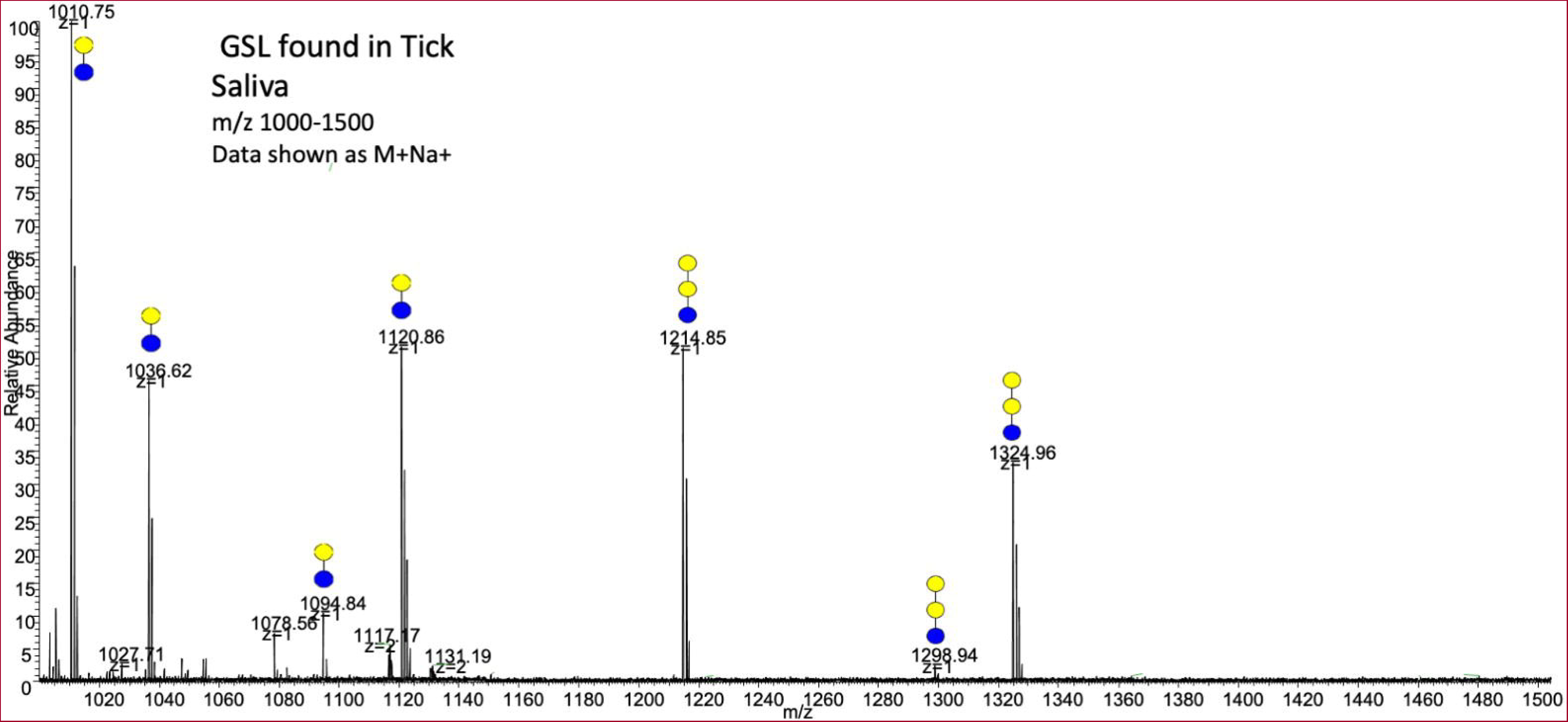
Mass spectrometry of tick saliva reveals evidences of alpha-gal bound lipids in saliva of partially fed *Am. americanum*.

### BAT Assay reveals basophil activation triggered by lipids isolated from tick saliva and salivary glands

Lipid profiling showed the presence of alpha-gal bound lipids in *Am. americanum* saliva, and salivary glands. Basophil activation testing was performed to investigate the putative role of alpha-gal conjugated salivary lipids. Basophils from a healthy, non-allergic donor were IgE-stripped and primed overnight with plasma from an individual with α-gal syndrome (α-gal sIgE = 53.7 IU/mL, total IgE = 74.8 IU/mL) as described earlier. The sensitized cells were exposed to different stimuli for 30 minutes, including Basophil media (Figure 2A), crosslinking anti-IgE antibody (positive control, Figure 2B), protein fractions lacking lipids from *Am. americanum* salivary gland (Figure 2C), lipid extracted from *Am. americanum* salivary gland (Figure 2D), protein fractions lacking lipids from *Am. americanum* saliva (Figure 2E), lipid extracted from partially fed saliva (Figure 2F). Flow cytometry assessed CD63 expression on lineage-1^-^HLA-DR^-^ CD123^+^CD203c^+^ basophils. The frequency of CD63^+^ basophils significantly increased following sensitization with α-gal allergic plasma and stimulation with α-gal-containing tick salivary lipid samples from *Am. americanum* (PF SG lipid extract from saliva and salivary gland, 74.1% and 71.2% respectively vs media/positive control) (p < 0.05 vs. media, Figure 2A, 2B, 2D, 2F). Protein fractions lacking lipids from tick salivary gland and saliva also induced increased basophil activation (49.3% and 60.7% respectively, Figure 2E, 2C) compared to control (1.26%, Figure 2A).

**Figure 2:**
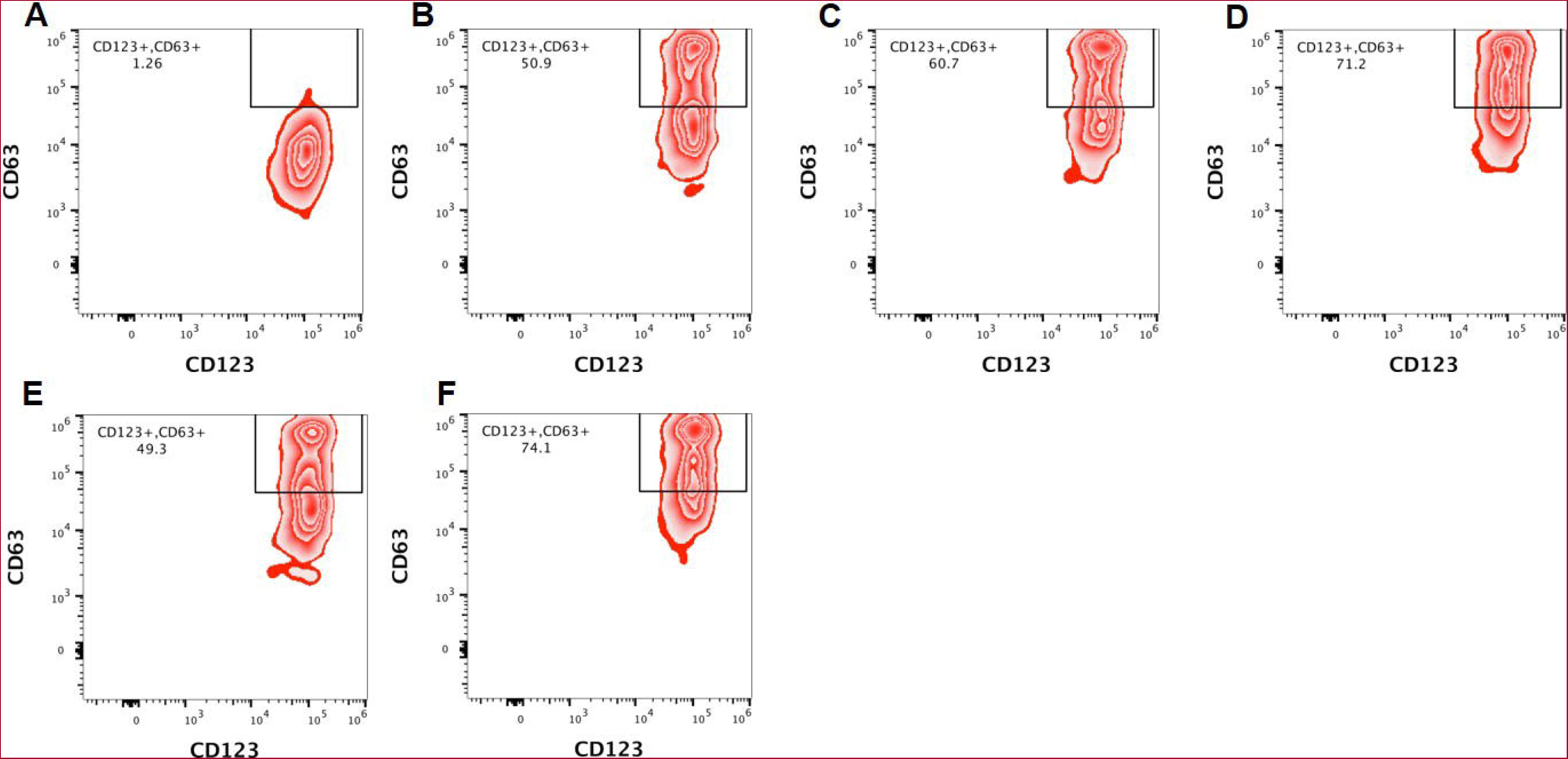
Flow cytometric analysis of human basophil activation by salivary total proteins of silenced tissues, lipids, and protein fractions of tick *Amblyomma Americanum (Am. Americanum)* saliva and salivary glands. Basophils from a healthy, non-allergic control subject, depleted of IgE, were overnight primed with plasma from an individual with α-gal syndrome (α-gal sIgE = 53.7 IU/ml, total IgE = 74.8 IU/ml). These sensitized cells were then exposed to various stimuli for 30 minutes, including **(A)** RPMI media, **(B)** crosslinking anti-IgE antibody (used as a positive control), **(C)** protein fraction left after extraction of lipid from partially blood feed *Am. americanum*. tick salivary gland, ***(D)*** lipid isolated from partially blood-feed *Am americanum* tick salivary glands, **(E)** protein fraction left after extraction of lipid from saliva, **(F)** lipid isolated from partially blood-feed *Am americanum*. tick saliva,. Flow cytometry was utilized to assess CD63 expression on lineage-1^-^HLA-DR^-^CD123^+^CD203c^+^ basophils. Results are representative of three biological replicates.

## Discussion

The Lone-star tick, originally limited to the Southeast, has now spread to include almost the entire U.S. east of the Rocky Mountains, transmitting diseases such as spotted fever group rickettsiosis, human monocytic ehrlichiosis, and tularemia (Paddock et al., 2003, Raghavan et al, 2019). Additionally, it is associated with delayed anaphylaxis to red meat (Commins et al., 2009). *Am. americanum*, the primary blood-feeding ectoparasite linked to causing food allergies in the U.S., has sparked significant attention. The discovery of α-gal in ticks and the development of α-gal-specific IgE in individuals with a history of tick bites play a pivotal role in allergic reactions to the α-gal antigen in red meat (Commins et al., 2009; Kerash et al., 2023).Tick saliva antigens are the key elements in AGS development (Crispell et al., 2019). Ticks bite humans and inject saliva, which delivers antigens containing α-gal epitopes triggering the production of anti-α-gal antibodies (Commins and Platts-Mills 2013). Recently, N-linked glycan analysis confirmed α-gal in the saliva and salivary glands of *Am. americanum* and *Ixodes scapularis* (Crispell et al., 2019). An indirect basophil activation test also confirmed the stimulation of high expression of CD63, a marker of human basophil degranulation by antigenic α-Gal of *Am. americanum* and *Ix. scapularis* ticks, but the functional role of *Ix. scapularis* α-gal in inducing IgE response is not known (Crispell et al., 2019). Although, the presence of α-gal conjugated saliva peptides has been characterized, the prevalence of lipid-bound α-gal in tick saliva has yet to be determined. Addressing this fundamental knowledge gap is important for solving the puzzle and enhancing our understanding of tick-induced food allergies (van Nunen et al., 2009; Crispell et al., 2019; Commins, 2020; Sharma and Karim, 2021; Cabezas-Cruz et al., 2018; Choudhary et al., 2021; Sharma et al., 2024).

Chaudhary et al, (2021) demonstrated that tick salivary gland extracts induced AGS in α-Gal^KO^ mice. Furthermore, including this study our recent study has established the fact that alpha-gal bound to protein is critical for AGS induction during host sensitization (Sharma et al, 2024). The study’s intriguing discovery indicates that not all hosts undergo AGS (Alpha-Gal Syndrome) induction after being bitten by a tick. This suggests the involvement of diverse factors associated with host sensitization. This observation is consistent with recent meta-analysis findings that examined clinical and experimental data regarding tick bites and AGS induction in individuals with a history of tick exposure. It is crucial to underscore that alpha-gal sensitization does not occur consistently after every tick bite, as highlighted by Young et al. (2021). This highlights the importance of both inherent factors in ticks and those linked to the host in this complex process, as noted by Sharma et al. (2021; 2024). A recently proposed hypothesis suggests that the initiation of AGS (Alpha-Gal Syndrome) development is associated with α-gal conjugated lipids, as outlined by studies conducted by Roman-Carrasco et al. (2021), Hopkins et al. (2022), and Chakrapani et al. (2022).

In this study, we aimed to verify the hypothesis concerning the identification of α-gal bound lipids implicated in hypersensitivity. The focus was on identifying alpha-gal lipid antigens potentially associated with the onset of Alpha-Gal Syndrome (AGS) following a tick bite. Through our NSI-MS analysis investigating glycan composition in partially fed *Am. americanum* saliva, we discovered the existence of lipids containing α-gal moieties, as detailed in Table 1. However, such lipids with α-gal moieties were not detected in the overall lipid composition extracted from partially blood-fed salivary glands (Table S1, Figures 1S, 2S, 3S). It is important to note that the profile of salivary gland was complex and multiple unknown signals were detected in NSI-MS spectera (Supplementary figure S1).

A mechanism of regulated exocytosis is involved in the secretion of saliva components, and saliva contains a plethora of bioactive molecules tha target a broad spectrum of host defense mechanisms to allow ticks to blood feed on the vertebrate host for several days (Karim et al., 2002). Identification of α-gal conjugated lipids in tick saliva may be due to lesser complexity of lipids and presence of the high concentration of lipids belonging to particular group in the saliva including Prostaglandin E2 in particular stage of tick feeding (Bowman et al., 1997). The complex lipid composition of tick salivary glands, which surpasses the complexity observed in saliva, presents a difficulty in verifying the existence of lipids containing attached glycan moieties. These lipids are identified as unknown peak signals in the NSI-MS analysis spectra data (refer to Supplementary Figures S1, S2, S3). The exact percentage of α-gal conjugated to lipids or peptides remains to be determined. However, identification of α-gal, even in trace amounts in saliva, supports the idea that lipid-bound α-gal may contribute to inducing hypersensitivity in humans along with α-gal conjugated salivary peptides. This is further supported by the activation of basophils by lipids extracted (deficient of α-gal conjugated peptides) from both saliva and salivary glands (Figure 2). Our initial study also indicated that the BAT assay, using the phospholipase-silenced salivary gland extract, resulted in a notable reduction in basophil activation (data not shown), a piece of correlative evidence showing the role of PLA2 in PGE2 synthesis (Qian et al., 1997). This finding underscores the crucial need for follow-up studies to determine the functional role of PLA2 in the synthesis of α-gal bound lipids, and their putative role in AGS development. Based on the observed size, α-gal bound lipids were discovered in tick saliva via NSI-MS, belonging to Gb3 or Isoglobotriosylceramide (iGb3Cer), a group of lipids (see Table 1), potentially consisting of carbon chains of 16, 18, 22, and 24. Previous research, focusing on the profiling of tick lipids following tick feeding, reported that lipids containing carbon chains of 16, 18, 22, and 24 corresponded to palmitic acid, oleic acid or octadecenoic acid, docosahexaenoic acid or DHA, and nervonic or cis-15-tetracosenoic acid, respectively (Shipley et al., 1993; Renthal et al., 2019).

Indeed, it was noted that *Am. americanum* may have putative enzymes responsible for the transfer of α-gal to lipids (Karim and Ribero, 2015). However, with current data, it is inconclusive whether these lipids are directly derived from host blood or synthesized by ticks using host precursor lipid molecules. Investigating how these alpha-gal structures accumulate in tick saliva poses an interesting aspect of future research.

Fundamentally, these tick salivary lipids were marked as important in tick physiology, energy, and reproduction (Renthal et al., 2019; Rees et al., 2004). A study focused on the analysis of lipid profiles following tick feeding revealed that the composition and abundance of salivary gland fatty acids change drastically during feeding. Notably, myristic (14:0) and palmitic acid (16:0) decrease following feeding; conversely, stearic acid (18:0) increases during feeding. Among all lipids, oleic acid (18:1) is the most abundant fatty acid throughout tick feeding (Shipley et al., 1993). The research findings from Renthal et al. (2019) reveal that salivary glands undergo notable alterations in mass and fatty acid composition linked to phospholipids during feeding. Specifically, the study identifies phosphatidylcholine (PC) and phosphatidylethanolamine (PE) as the primary phospholipids present, while triglycerides (TGs) emerge as the predominant neutral lipids within the glands.

In the context of lipid abundantly attached to α-gal, it is reported that α-gal is predominantly linked to glycolipids. Glycolipids are a type of lipid molecule containing a carbohydrate (glycan) moiety covalently attached to a lipid moiety. The source of these glycolipids in ticks may be derived from host blood because ticks strictly rely on host blood for energy (Alsmari and Wall, 2020). In the context of tick saliva, glycolipids can be present and play various roles in tick-host interaction. The composition of saliva in ticks is complex and involves a mixture of different molecules, including proteins, lipids, and carbohydrates (Neelakanta & Sultana, 2022). It’s important to note that the detailed characterization of specific lipids and the glycosylation pattern in lipids present in tick saliva, their structures, and their functions would require advanced analytical techniques, such as mass spectrometry or chromatography, coupled with immunological or biochemical assays. Researchers studying tick biology and tick-host interactions may investigate the presence and roles of glycolipids as part of their broader exploration of tick saliva composition and function.

Humans without α-gal hypersensitivity typically exhibit up to 1% of circulating IgG antibodies specific to anti-gal, signifying the immune system’s recognition of this carbohydrate antigen. Considering ticks’ extended attachment to hosts, the continuous secretion of small amounts of α-gal and other antigenic molecules could prompt an immune response, resulting in the development of an IgE response against α-gal. Our findings signify fresh perspectives on tick physiology and support the hypothesis of hypersensitivity reactions initiated post-parasitism by ticks. It advances our fundamental knowledge and comprehension of how ticks acquire and transmit pathogenic α-gal antigens to the host, potentially offering avenues for future medical condition treatment or prevention. Notably, the research reveals that tick saliva contains alpha-gal-bound lipid markers, and basophil activation tests demonstrate these lipid antigens’ potential to activate basophils. Key questions about how these lipid antigens facilitate host sensitization post-tick bites remain to be explored. From these observations, it can be inferred that certain ticks, like *Am. americanum*, possess a unique ability to induce Alpha-Gal Syndrome (AGS) after a tick bite. This capability is attributed to the intrinsic tick machinery associated with the development of the alpha-gal antigen during blood feeding.

Our findings establish 1) *Am. americanum* tick saliva contains alpha-gal-bound lipid antigens, 2) the basophil activation test establishes the ability of isolated alpha-gal-bound lipids and protein to activate basophils, implying their potential as antigens capable of sensitizing the host during a tick bite. These results provide novel insights into understanding the role of essential intrinsic factors in ticks, particularly the presence of lipid-bound alpha-gal antigens potentially involved in sensitizing the host during hematophagy.

## Author Contributions

Conceptualization: Surendra Raj Sharma, Shahid Karim

Methodology: Surendra Raj Sharma, Sabir Hussain Shailesh Choudhary, Commins, Shahid Karim

Data curation: Surendra Raj Sharma, Shailesh Choudhary, Scott Commins, Shahid Karim

Funding acquisition: Scott Commins, Shahid Karim

Investigation: Surendra Sharma, Shailesh Choudhary, Scott Commins, Shahid Karim

Project administration: Shahid Karim

Resources: Scott Commins, Shahid Karim

Supervision; Shahid Karim

Writing, original draft: Surendra Raj Sharma, Shailesh Choudhary, Scott Commins, Shahid Karim

Writing, review & editing: Surendra Sharma, Shailesh Choudhary, Scott Commins, Shahid Karim

All authors read and approved the manuscript.

## References

Adegoke, A., Kumar, D., Bobo, C., Rashid, M. I., Durrani, A. Z., Sajid, M. S., & Karim, S. (2020). Tick-Borne Pathogens Shape the Native Microbiome Within Tick Vectors. Microorganisms, 8(9), 1299. 10.3390/microorganisms8091299

Apostolovic, D., Tran, T. A., Hamsten, C., Starkhammar, M., Cirkovic Velickovic, T., & van Hage, M. (2014). Immunoproteomics of processed beef proteins reveal novel galactose-α-1,3-galactose-containing allergens. Allergy, 69(10), 1308–1315. 10.1111/all.12462

Berg, E. A., Platts-Mills, T. A., & Commins, S. P. (2014). Drug allergens and food--the cetuximab and galactose-α-1,3-galactose story. Annals of allergy, asthma & immunology: official publication of the American College of Allergy, Asthma, & Immunology, 112(2), 97–101. 10.1016/j.anai.2013.11.014

Binder, A. M., Commins, S. P., Altrich, M. L., Wachs, T., Biggerstaff, B. J., Beard, C. B., Petersen, L. R., Kersh, G. J., & Armstrong, P. A. (2021). Diagnostic testing for galactose-alpha-1,3-galactose, United States, 2010 to 2018. Annals of allergy, asthma & immunology: official publication of the American College of Allergy, Asthma, & Immunology, 126(4), 411–416.e1.

Bullard, R., Sharma, S. R., Das, P. K., Morgan, S. E., & Karim, S. (2019). Repurposing of Glycine-Rich Proteins in Abiotic and Biotic Stresses in the Lone-Star Tick (Amblyomma americanum). Frontiers in physiology, 10, 744. 10.3389/fphys.2019.00744

Román-Carrasco, P., Hemmer, W., Cabezas-Cruz, A., Hodžić, A., de la Fuente, J., & Swoboda, I. (2021). The α-Gal Syndrome and Potential Mechanisms. Frontiers in allergy, 2, 783279. 10.3389/falgy.2021.783279

Chakrapani, N., Fischer, J., Swiontek, K., Codreanu-Morel, F., Hannachi, F., Morisset, M., Mugemana, C., Bulaev, D., Blank, S., Bindslev-Jensen, C., Biedermann, T., Ollert, M., & Hilger, C. (2022). α-Gal present on both glycolipids and glycoproteins contributes to immune response in meat-allergic patients. The Journal of allergy and clinical immunology, 150(2), 396–405.e11. 10.1016/j.jaci.2022.02.030

Chmelař, J., Kotál, J., Kopecký, J., Pedra, J. H. F., & Kotsyfakis, M. (2016). All For One and One For All on the Tick-Host Battlefield. Trends in parasitology, 32(5), 368–377. 10.1016/j.pt.2016.01.004

Choudhary, S. K., Karim, S., Iweala, O. I., Choudhary, S., Crispell, G., Sharma, S. R., Addison, C. T., Kulis, M., Herrin, B. H., Little, S. E., & Commins, S. P. (2021). Tick salivary gland extract induces alpha-gal syndrome in alpha-gal deficient mice. Immunity, inflammation and disease, 9(3), 984–990. 10.1002/iid3.457

Commins, S. P., James, H. R., Kelly, L. A., Pochan, S. L., Workman, L. J., Perzanowski, M. S., Kocan, K. M., Fahy, J. V., Nganga, L. W., Ronmark, E., Cooper, P. J., & Platts-Mills, T. A. (2011). The relevance of tick bites to the production of IgE antibodies to the mammalian oligosaccharide galactose-α-1,3-galactose. The Journal of allergy and clinical immunology, 127(5), 1286–93.e6. 10.1016/j.jaci.2011.02.019

Commins, S. P., James, H. R., Stevens, W., Pochan, S. L., Land, M. H., King, C., Mozzicato, S., & Platts-Mills, T. A. (2014). Delayed clinical and ex vivo response to mammalian meat in patients with IgE to galactose-alpha-1,3-galactose. The Journal of allergy and clinical immunology, 134(1), 108–115. 10.1016/j.jaci.2014.01.024

Commins, S. P., Satinover, S. M., Hosen, J., Mozena, J., Borish, L., Lewis, B. D., Woodfolk, J. A., & Platts-Mills, T. A. (2009). Delayed anaphylaxis, angioedema, or urticaria after consumption of red meat in patients with IgE antibodies specific for galactose-alpha-1,3-galactose. The Journal of allergy and clinical immunology, 123(2), 426–433. 10.1016/j.jaci.2008.10.052

Crispell, G., Commins, S. P., Archer-Hartman, S. A., Choudhary, S., Dharmarajan, G., Azadi, P., & Karim, S. (2019). Discovery of Alpha-Gal-Containing Antigens in North American Tick Species Believed to Induce Red Meat Allergy. Frontiers in immunology, 10, 1056. 10.3389/fimmu.2019.01056

Fischer, J., Hebsaker, J., Caponetto, P., Platts-Mills, T. A., & Biedermann, T. (2014). Galactose-alpha-1,3-galactose sensitization is a prerequisite for pork-kidney allergy and cofactor-related mammalian meat anaphylaxis. The Journal of allergy and clinical immunology, 134(3), 755–759.e1. 10.1016/j.jaci.2014.05.051

Galili U. (1999). Evolution of alpha 1,3galactosyltransferase and of the alpha-Gal epitope. Sub-cellular biochemistry, 32, 1–23. 10.1007/978-1-4615-4771-6_1

Galili, U., Avila, J. L. (Eds.) (1999). α-Gal and Anti-Gal, α1,3-Galactosyltransferase, α-Gal Epitopes, and the Natural Anti-Gal Antibody Subcellular Biochemistry Vol. 32 (Boston, MA: Springer US). doi: 10.1007/978-1-4615-4771-6

Hamsten, C., Starkhammar, M., Tran, T. A., Johansson, M., Bengtsson, U., Ahlén, G., Sällberg, M., Grönlund, H., & van Hage, M. (2013). Identification of galactose-α-1,3-galactose in the gastrointestinal tract of the tick Ixodes ricinus; possible relationship with red meat allergy. Allergy, 68(4), 549–552. 10.1111/all.12128

Hopkins, G. V., Cochrane, S., Onion, D., & Fairclough, L. C. (2022). The Role of Lipids in Allergic Sensitization: A Systematic Review. Frontiers in molecular biosciences, 9, 832330. 10.3389/fmolb.2022.832330 10.1016/j.anai.2020.12.019

Iweala, O. I., Choudhary, S. K., Addison, C. T., Batty, C. J., Kapita, C. M., Amelio, C., Schuyler, A. J., Deng, S., Bachelder, E. M., Ainslie, K. M., Savage, P. B., Brennan, P. J., & Commins, S. P. (2020). Glycolipid-mediated basophil activation in alpha-gal allergy. The Journal of allergy and clinical immunology, 146(2), 450–452. 10.1016/j.jaci.2020.02.006

Karim, S., Singh, P., & Ribeiro, J. M. (2011). A deep insight into the sialotranscriptome of the gulf coast tick, Amblyomma maculatum. PloS one, 6(12), e28525. 10.1371/journal.pone.0028525

Kersh, G. J., Salzer, J., Jones, E. S., Binder, A. M., Armstrong, P. A., Choudhary, S. K., Commins, G. K., Amelio, C. L., Kato, C. Y., Singleton, J., Biggerstaff, B. J., Beard, C. B., Petersen, L. R., & Commins, S. P. (2023). Tick bite as a risk factor for alpha-gal-specific immunoglobulin E antibodies and development of alpha-gal syndrome. Annals of allergy, asthma & immunology: official publication of the American College of Allergy, Asthma, & Immunology, 130(4), 472–478. 10.1016/j.anai.2022.11.021

Macher, B. A., & Galili, U. (2008). The Galalpha1,3Galbeta1,4GlcNAc-R (alpha-Gal) epitope: a carbohydrate of unique evolution and clinical relevance. Biochimica et biophysica acta, 1780(2), 75–88. 10.1016/j.bbagen.2007.11.003

Monzón, J. D., Atkinson, E. G., Henn, B. M., & Benach, J. L. (2016). Population and Evolutionary Genomics of Amblyomma americanum, an Expanding Arthropod Disease Vector. Genome biology and evolution, 8(5), 1351–1360. 10.1093/gbe/evw080

Patrick, C. D., & Hair, J. A. (1975). Laboratory rearing procedures and equipment for multi-host ticks (Acarina: Ixodidae). Journal of medical entomology, 12(3), 389–390. 10.1093/jmedent/12.3.389

Paddock, C. D., & Childs, J. E. (2003). Ehrlichia chaffeensis: a prototypical emerging pathogen. Clinical microbiology reviews, 16(1), 37–64. 10.1128/CMR.16.1.37-64.2003

Raghavan, R. K., Peterson, A. T., Cobos, M. E., Ganta, R., & Foley, D. (2019). Current and Future Distribution of the Lone Star Tick, Amblyomma americanum (L.) (Acari: Ixodidae) in North America. PloS one, 14(1), e0209082. 10.1371/journal.pone.0209082

Roseman S. (2001). Reflections on glycobiology. The Journal of biological chemistry, 276(45), 41527–41542. 10.1074/jbc.R100053200

Sharma, S. R., & Karim, S. (2021). Tick Saliva and the Alpha-Gal Syndrome: Finding a Needle in a Haystack. Frontiers in cellular and infection microbiology, 11, 680264. 10.3389/fcimb.2021.680264

Takahashi, H., Chinuki, Y., Tanaka, A., & Morita, E. (2014). Laminin γ-1 and collagen α-1 (VI) chain are galactose-α-1,3-galactose-bound allergens in beef. Allergy, 69(2), 199–207. 10.1111/all.12302

Thompson, J. M., Carpenter, A., Kersh, G. J., Wachs, T., Commins, S. P., & Salzer, J. S. (2023). Geographic Distribution of Suspected Alpha-gal Syndrome Cases - United States, January 2017-December 2022. MMWR. Morbidity and mortality weekly report, 72(30), 815–820. 10.15585/mmwr.mm7230a2

van Nunen, S. A., O’Connor, K. S., Clarke, L. R., Boyle, R. X., & Fernando, S. L. (2009). An association between tick bite reactions and red meat allergy in humans. The Medical journal of Australia, 190(9), 510–511. 10.5694/j.1326-5377.2009.tb02533.x

Shipley, M. M., Dillwith, J. W., Bowman, A. S., Essenberg, R. C., & Sauer, J. R. (1993). Changes in lipids of the salivary glands of the lone star tick, Amblyomma americanum, during feeding. The Journal of parasitology, 79(6), 834–842.

Renthal, R., Lohmeyer, K., Borges, L. M. F., & Pérez de León, A. A. (2019). Surface lipidome of the lone star tick, Amblyomma americanum, provides leads on semiochemicals and lipid metabolism. Ticks and tick-borne diseases, 10(1), 138–145. 10.1016/j.ttbdis.2018.09.009

Rees H. H. (2004). Hormonal control of tick development and reproduction. Parasitology, 129 Suppl, S127–S143. 10.1017/s003118200400530x

Karim, S., & Ribeiro, J. M. (2015). An Insight into the Sialome of the Lone Star Tick, Amblyomma americanum, with a Glimpse on Its Time Dependent Gene Expression. PloS one, 10(7), e0131292. 10.1371/journal.pone.0131292

Alasmari, S., & Wall, R. (2020). Determining the total energy budget of the tick Ixodes ricinus. Experimental & applied acarology, 80(4), 531–541. 10.1007/s10493-020-00479-1

Neelakanta, G., & Sultana, H. (2022). Tick Saliva and Salivary Glands: What Do We Know So Far on Their Role in Arthropod Blood Feeding and Pathogen Transmission. Frontiers in cellular and infection microbiology, 11, 816547. 10.3389/fcimb.2021.816547

Qian, Y., Essenberg, R. C., Dillwith, J. W., Bowman, A. S., & Sauer, J. R. (1997). A specific prostaglandin E2 receptor and its role in modulating salivary secretion in the female tick, Amblyomma americanum (L.). Insect biochemistry and molecular biology, 27(5), 387–395. 10.1016/s0965-1748(97)00010-6

