## Supplementary material for "Identification of Alpha-Gal conjugated Lipids in Saliva of Lone-Star Tick (*Amblyomma americanum*)": Fig. S1-S4, Table S1

### Slide 1
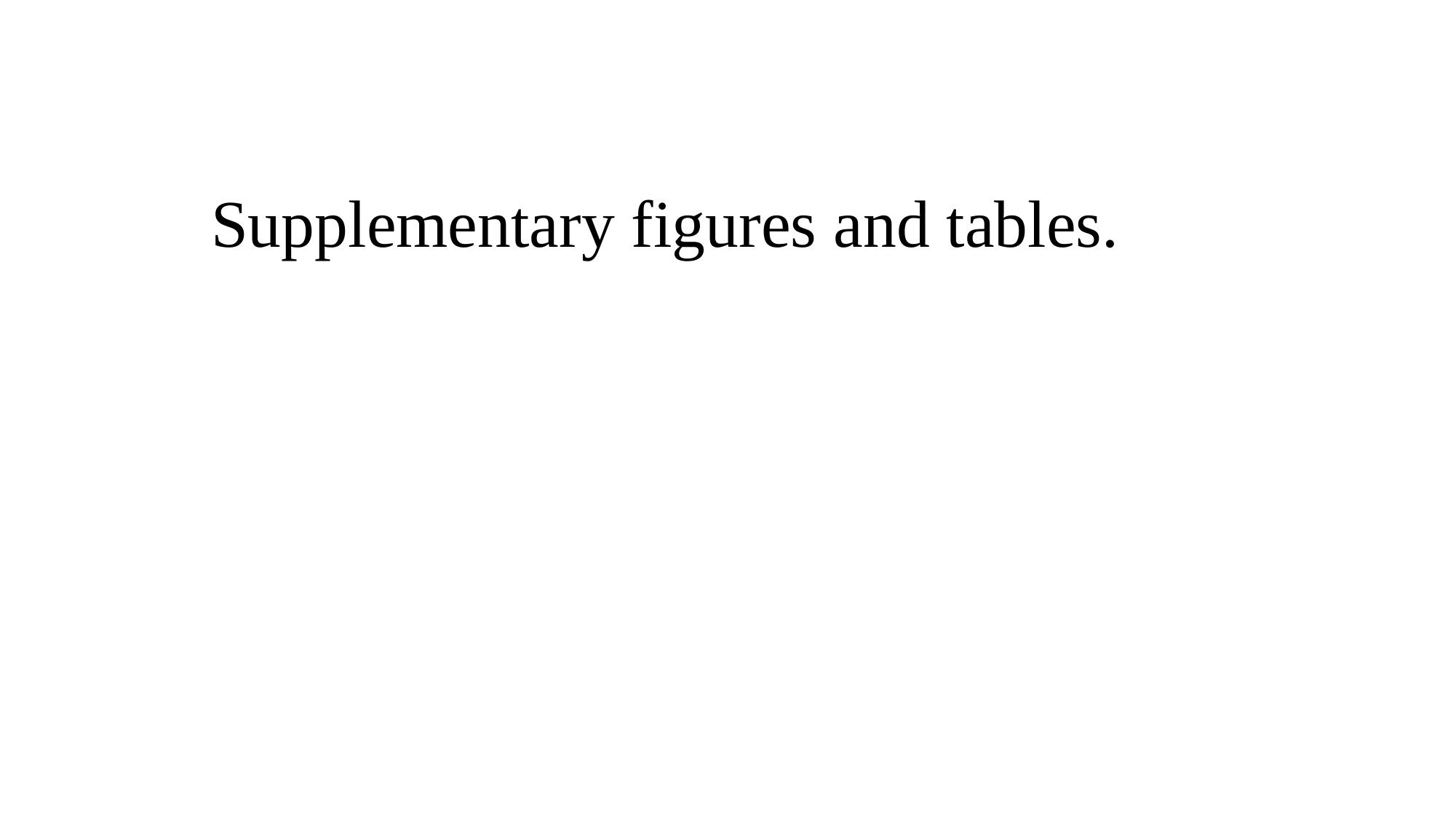

#### Supplementary figures and tables.

### Slide 2
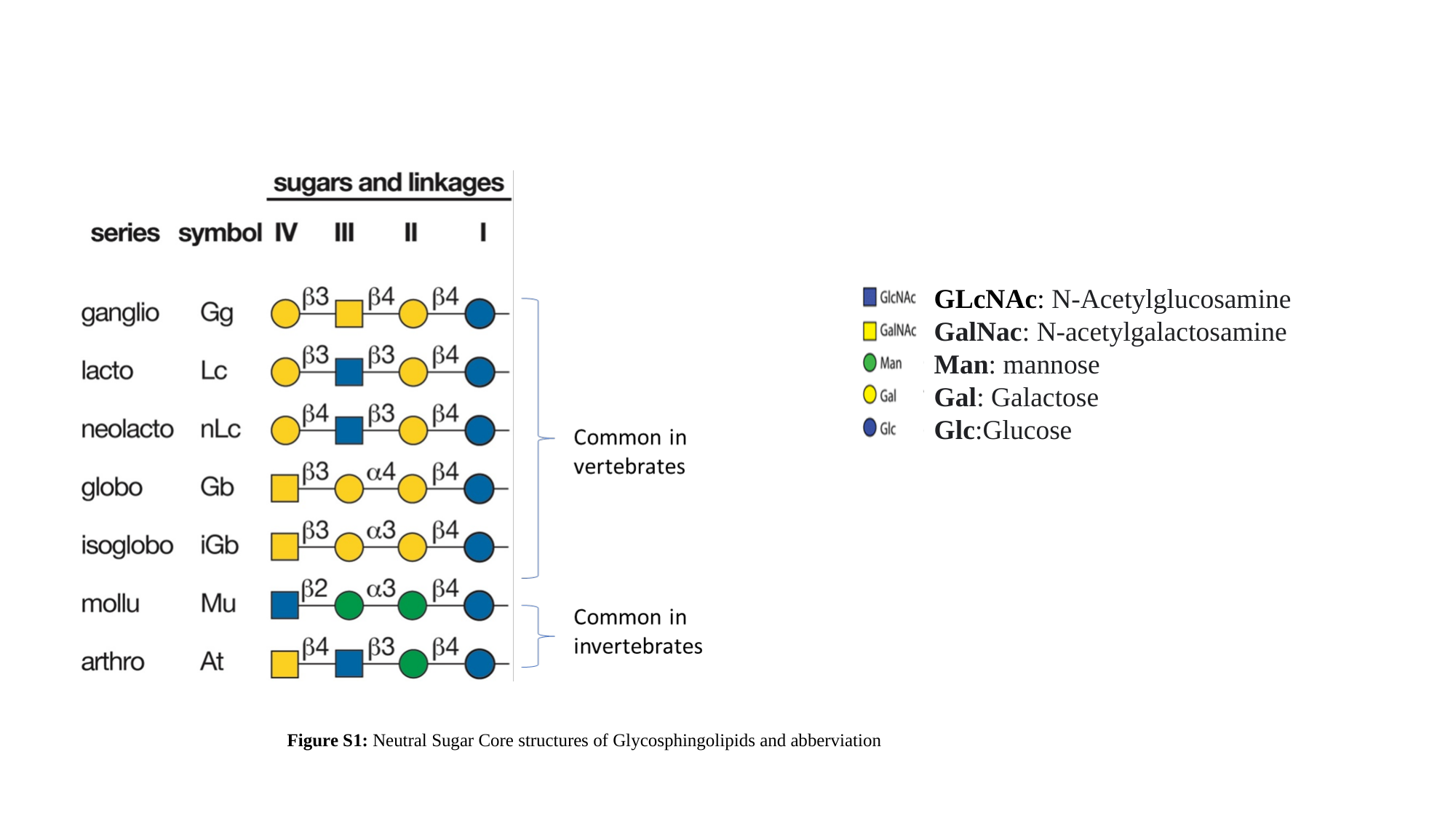

GLcNAc: N-Acetylglucosamine
GalNac: N-acetylgalactosamine
Man: mannose
Gal: Galactose
Glc:Glucose
Figure S1: Neutral Sugar Core structures of Glycosphingolipids and abberviation

### Slide 3
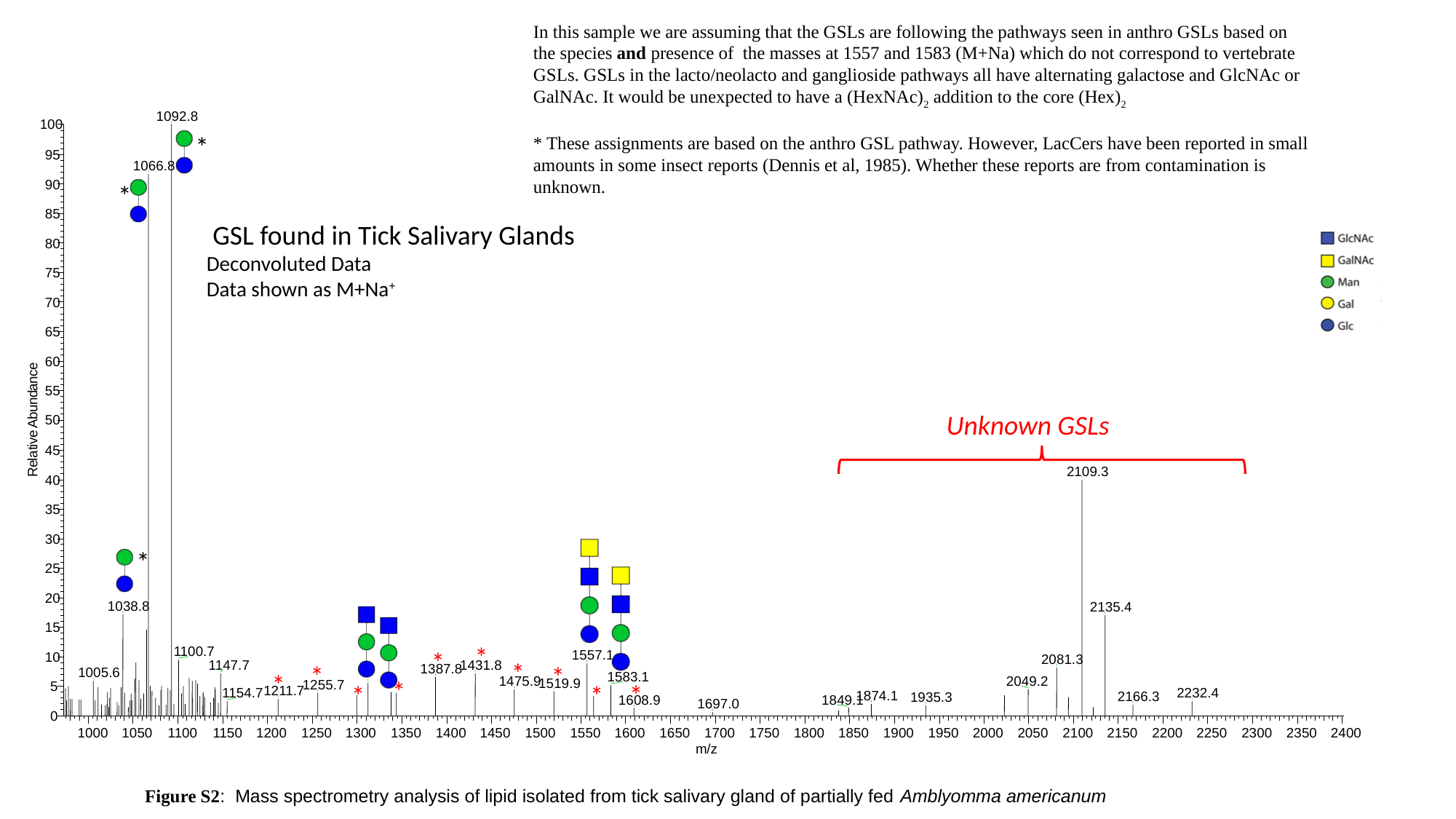

In this sample we are assuming that the GSLs are following the pathways seen in anthro GSLs based on the species and presence of  the masses at 1557 and 1583 (M+Na) which do not correspond to vertebrate GSLs. GSLs in the lacto/neolacto and ganglioside pathways all have alternating galactose and GlcNAc or GalNAc. It would be unexpected to have a (HexNAc)_2_ addition to the core (Hex)_2_
* These assignments are based on the anthro GSL pathway. However, LacCers have been reported in small amounts in some insect reports (Dennis et al, 1985). Whether these reports are from contamination is unknown.
1092.8
100
95
1066.8
90
85
80
75
70
65
60
e
c
n
a
55
d
n
u
b
50
A
e
v
i
t
45
a
l
e
R
2109.3
40
35
30
25
20
1038.8
2135.4
15
1100.7
1557.1
10
2081.3
1147.7
1431.8
1387.8
1005.6
1583.1
1475.9
2049.2
1519.9
1255.7
5
1211.7
1154.7
2232.4
1874.1
2166.3
1935.3
1608.9
1849.1
1697.0
0
1000
1050
1100
1150
1200
1250
1300
1350
1400
1450
1500
1550
1600
1650
1700
1750
1800
1850
1900
1950
2000
2050
2100
2150
2200
2250
2300
2350
2400
m/z
*
*
 GSL found in Tick Salivary Glands
Deconvoluted Data
Data shown as M+Na^+^
*
*
*
*
*
*
*
*
*
*
*
Unknown GSLs
Figure S2:  Mass spectrometry analysis of lipid isolated from tick salivary gland of partially fed Amblyomma americanum ​

### Slide 4
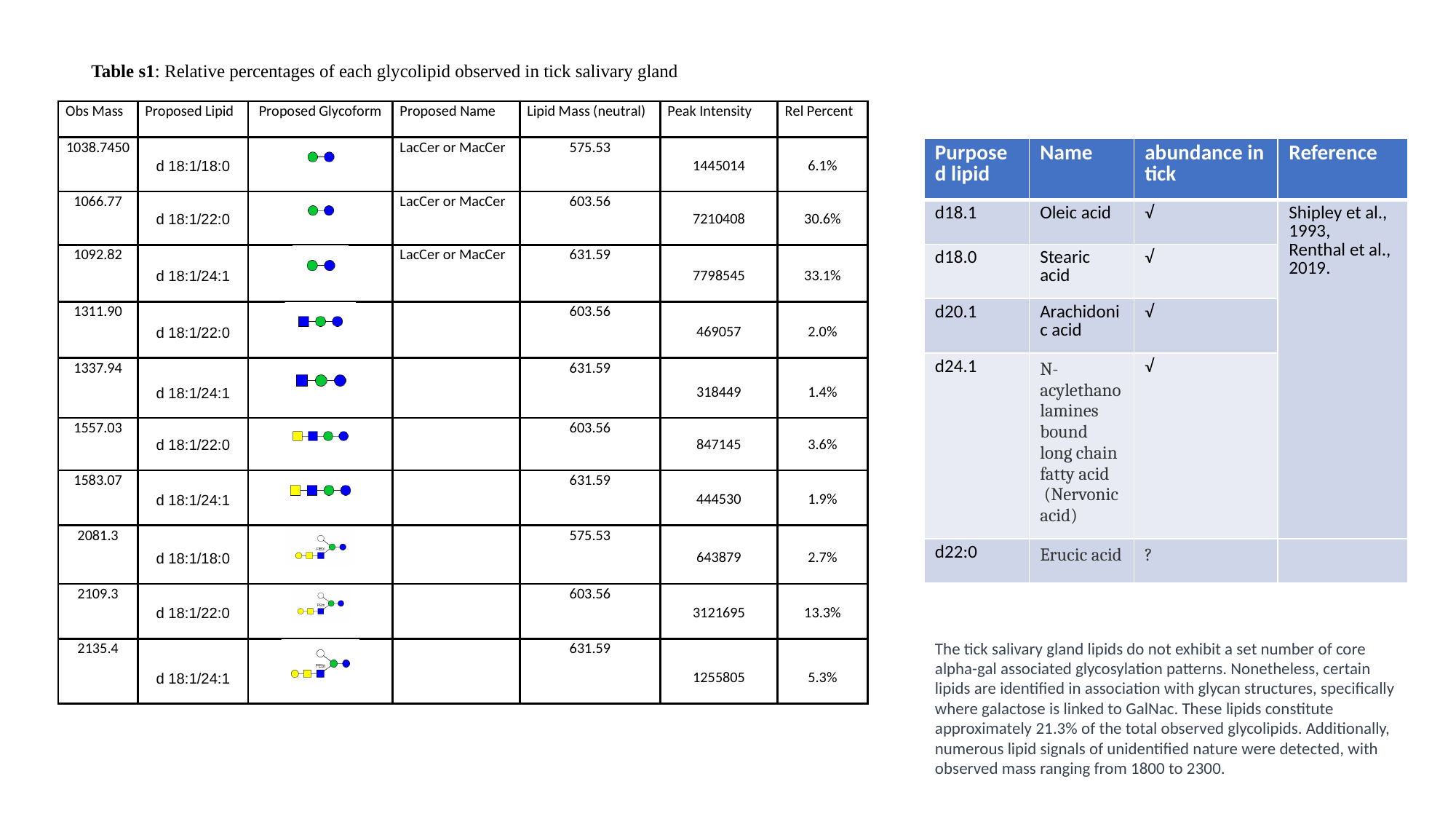

Table s1: Relative percentages of each glycolipid observed in tick salivary gland
| Purposed lipid | Name | abundance in tick | Reference |
| --- | --- | --- | --- |
| d18.1 | Oleic acid | √ | Shipley et al., 1993, Renthal et al., 2019. |
| d18.0 | Stearic acid | √ | |
| d20.1 | Arachidonic acid | √ | |
| d24.1 | N-acylethanolamines bound long chain fatty acid   (Nervonic acid) | √ | |
| d22:0 | Erucic acid | ? | |
The tick salivary gland lipids do not exhibit a set number of core alpha-gal associated glycosylation patterns. Nonetheless, certain lipids are identified in association with glycan structures, specifically where galactose is linked to GalNac. These lipids constitute approximately 21.3% of the total observed glycolipids. Additionally, numerous lipid signals of unidentified nature were detected, with observed mass ranging from 1800 to 2300.

### Slide 5
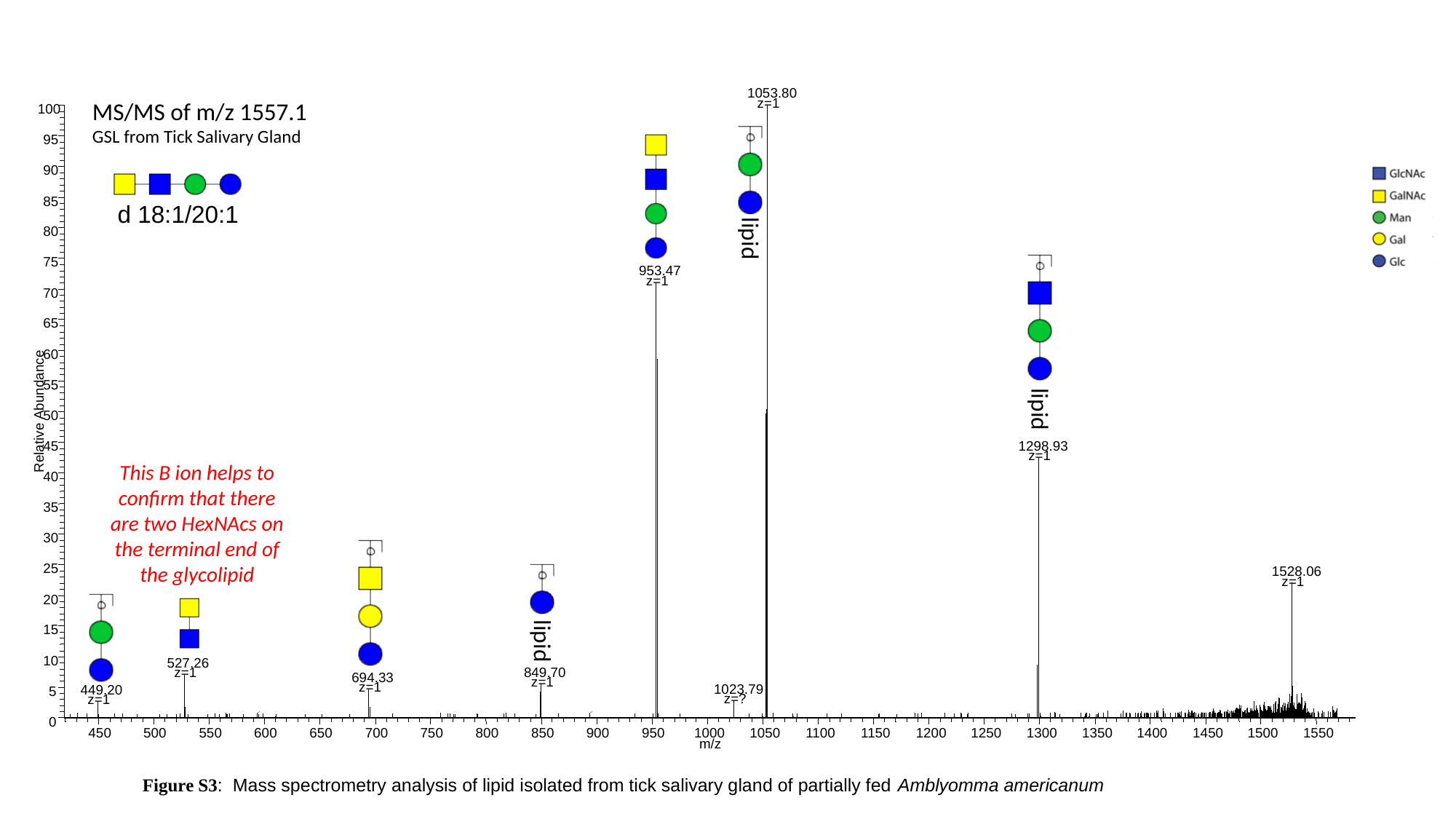

1053.80
MS/MS of m/z 1557.1
GSL from Tick Salivary Gland
z=1
100
95
90
85
d 18:1/20:1
lipid
80
75
953.47
z=1
70
65
60
55
lipid
Relative Abundance
50
45
1298.93
z=1
40
35
30
25
1528.06
z=1
20
15
lipid
10
527.26
z=1
849.70
694.33
z=1
z=1
1023.79
449.20
5
z=?
z=1
0
450
500
550
600
650
700
750
800
850
900
950
1000
1050
1100
1150
1200
1250
1300
1350
1400
1450
1500
1550
m/z
This B ion helps to confirm that there are two HexNAcs on the terminal end of the glycolipid
Figure S3:  Mass spectrometry analysis of lipid isolated from tick salivary gland of partially fed Amblyomma americanum ​

### Slide 6
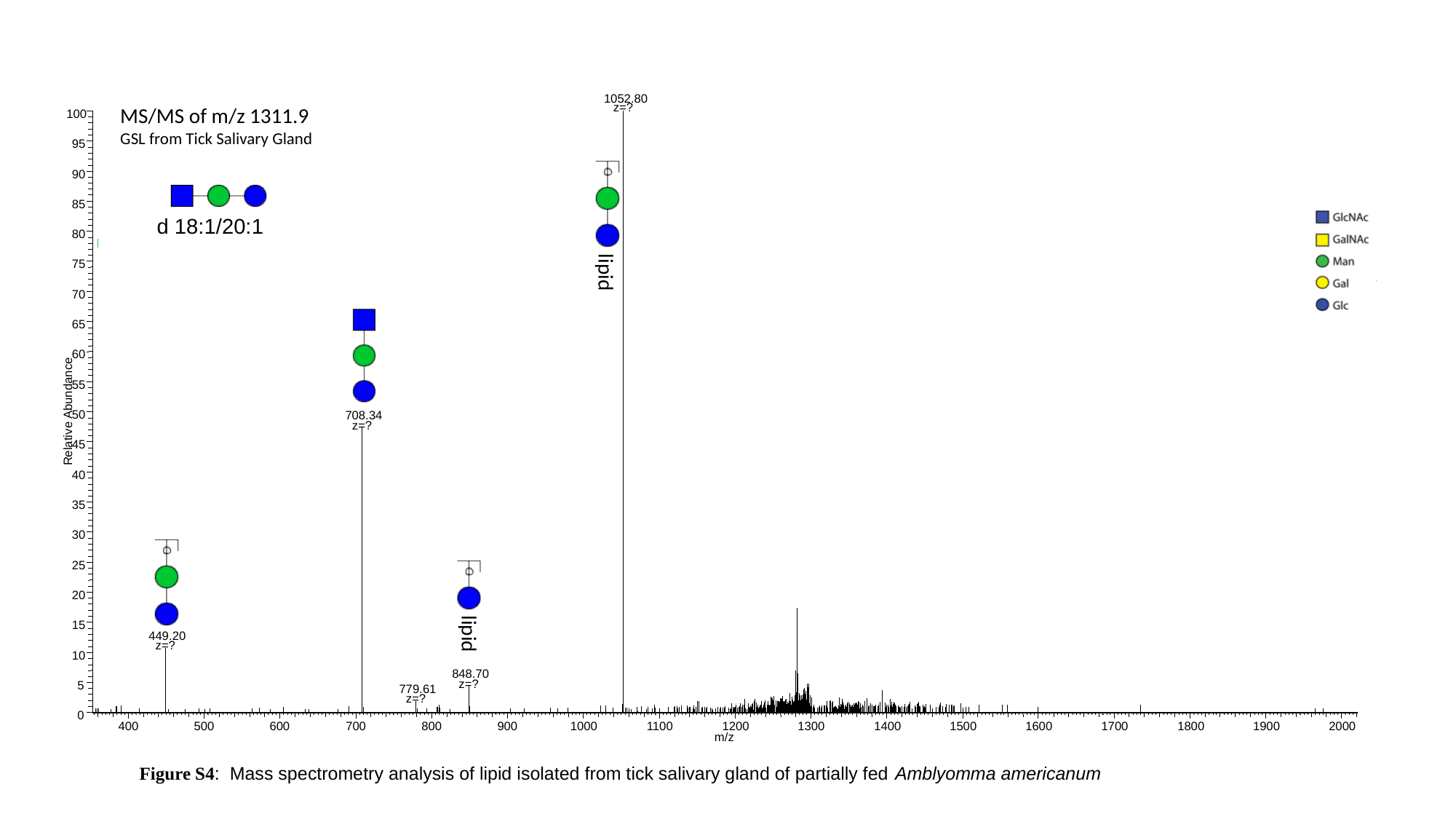

1052.80
MS/MS of m/z 1311.9
GSL from Tick Salivary Gland
z=?
100
95
90
85
d 18:1/20:1
80
75
lipid
70
65
60
55
Relative Abundance
50
708.34
z=?
45
40
35
30
25
20
15
lipid
449.20
z=?
10
848.70
z=?
5
779.61
z=?
0
400
500
600
700
800
900
1000
1100
1200
1300
1400
1500
1600
1700
1800
1900
2000
m/z
Figure S4:  Mass spectrometry analysis of lipid isolated from tick salivary gland of partially fed Amblyomma americanum ​
